# Junction-skipping regulation in complex disease

**DOI:** 10.1101/804039

**Authors:** Ruize Liu, Juha Karjalainen, Andrea Byrnes, Beryl B. Cummings, Padhraig Gormley, Yunfeng Ruan, Yongjin Park, Lei Hou, Khoi T. Nguyen, Ramnik Xavier, Mark J. Daly, Hailiang Huang

## Abstract

Multiple mRNA isoforms can be generated from a single gene locus through alternative splicing. Abnormality in alternative splicing has been linked to many human disorders. Here using RNA-seq data from 48 tissues from GTEx v7 release and summary statistics from GWAS of complex diseases and traits, we present a study to identify genomic variants regulating junction-skipping with the goal to understand their contribution to complex diseases and traits. For each tissue, we found 48 - 575 junction-skipping events regulated by genomic variants. We performed fine-mapping on both the junction-skipping association and 23 complex disease and trait associations and mapped them to 95% credible sets. We found 13 - 279 junction-skipping regulations were mapped to a credible set with ≤5 variants. On the genome-wide scale, we noted a clear disease-tissue specificity. Results from this approach provided critical insights into the functional mechanism of the genetic disease associations and contributed to our understanding of the genetic architecture of human complex disorders.

## Introduction

A single genetic locus can be transcribed to multiple RNA transcript isoforms due to alternative splicing^1^. Different isoforms transcribed from the same genetic locus can be translated to peptides with various structures and functions^2^. Many alternative splicing events are regulated in a cell-type or tissue-specific manner, at different developmental stages and disease processes. Alternative splicing mechanisms exist widely: the number of known mRNA transcripts are more than ten times the number of unique genes in the genome^3^. Therefore, alternative splicing increases the complexity of gene expression and plays an important role in diverse cellular processes and organism development.

Genome-wide association studies (GWAS) have identified a large number of variants associated with diseases. Most of these disease-associated variants are in the noncoding genome such as intronic and intergenic regions^4^. Understanding the molecular function of these disease-associated non-coding variants is a major challenge in the post-GWAS era^5^. Previous studies have shown that the vast majority of these variants might affect protein expression through regulating the transcription, splicing, or mRNA stability^3,6^. Intriguingly, disease-associated variants appear to be enriched in regulatory elements active in disease-associated cell types^5^.

Studies have shown that alternative splicing can be regulated by genomic variants^7^, and these splicing regulatory variants are major contributors to complex diseases^8^, potentially through the production of aberrant proteins^9^. Understanding the connection between the splicing regulatory variants and disease-associated variants will help to interpret the GWAS findings and provide insights into the disease pathogenesis.

Previously, alternative splicing has been characterized using the transcripted-focused or the exon/intron-focused methods using RNA sequencing (RNA-seq) data. The transcript-focused methods model the splicing through transcript isoform levels^10^, which requires the inference of full-length mRNA isoforms from short reads. These methods are therefore sensitive to isoform annotations^11^ and can be less accurate if the read depth is limited. The exon/intron-focused methods model the splicing through the inclusion/exclusion of a single or multiple exons/introns^12–15^. These methods do not require the prior knowledge of the isoforms, and have been shown to have better power due to its reduced complexity^14^.

Here we proposed a novel approach to characterize the alternative splicing with further reduced complexity and increased power by focusing on the intron-exon junctions. We note that many pathogenic transcript isoforms were generated by disruptions in the intron-exon junctions ^16–18^. By inferring the junction-skipping level directly using RNA-seq reads covering the junctions, we achieve a better accuracy and more power to capture such pathogenic alternative splicing events^14,15^.

We further identified variants regulating the junction-skipping rates (jsQTL) and investigated their connections with human complex disorders. We made two major improvements upon existing studies: 1) we improved the power to identify a jsQTL by applying a homogeneity test on the splicing pattern across individuals. By only focusing on splicing events that are heterogeneous across individuals, we reduced the multiple testing burden and increased the statistical power; 2) we performed fine-mapping to precisely map both the jsQTL and the disease-associated variants, which improves the signal-to-noise ratio and leads to better sensitivity. We applied this method to 48 tissues and cell lines in GTEx v7 release^19^ (Supplementary Table S1) and to 23 human complex traits/disorders (Supplementary Table S2), providing insights into the contribution of jsQTL to human complex traits/disorders in a tissue-specific manner.

## Results

We designed a method to identify junction-skipping events from the RNA-seq data from GTEx and their regulatory variants (Figure 1). To do this, we pooled the uniquely mapped junction reads in samples from the same tissue to identify junction-skipping events. This was done for all intron-exon junctions in the canonical transcripts. 48 tissues from GTEx were included in this study, with 80 - 491 samples per tissue (Supplementary Table S1). For a tissue and a junction of interest, we counted the number of reads supported this junction in the canonical transcript in individual *i* as *n*_*i*_, and the number of reads that skipped this junction as *s*_*i*_. We retained the junction if both 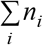 and 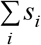 are greater than zero. The number of junctions included in the analysis ranged from 60,450 (substantia nigra) to 103,147 (lung).

**Figure 1.**
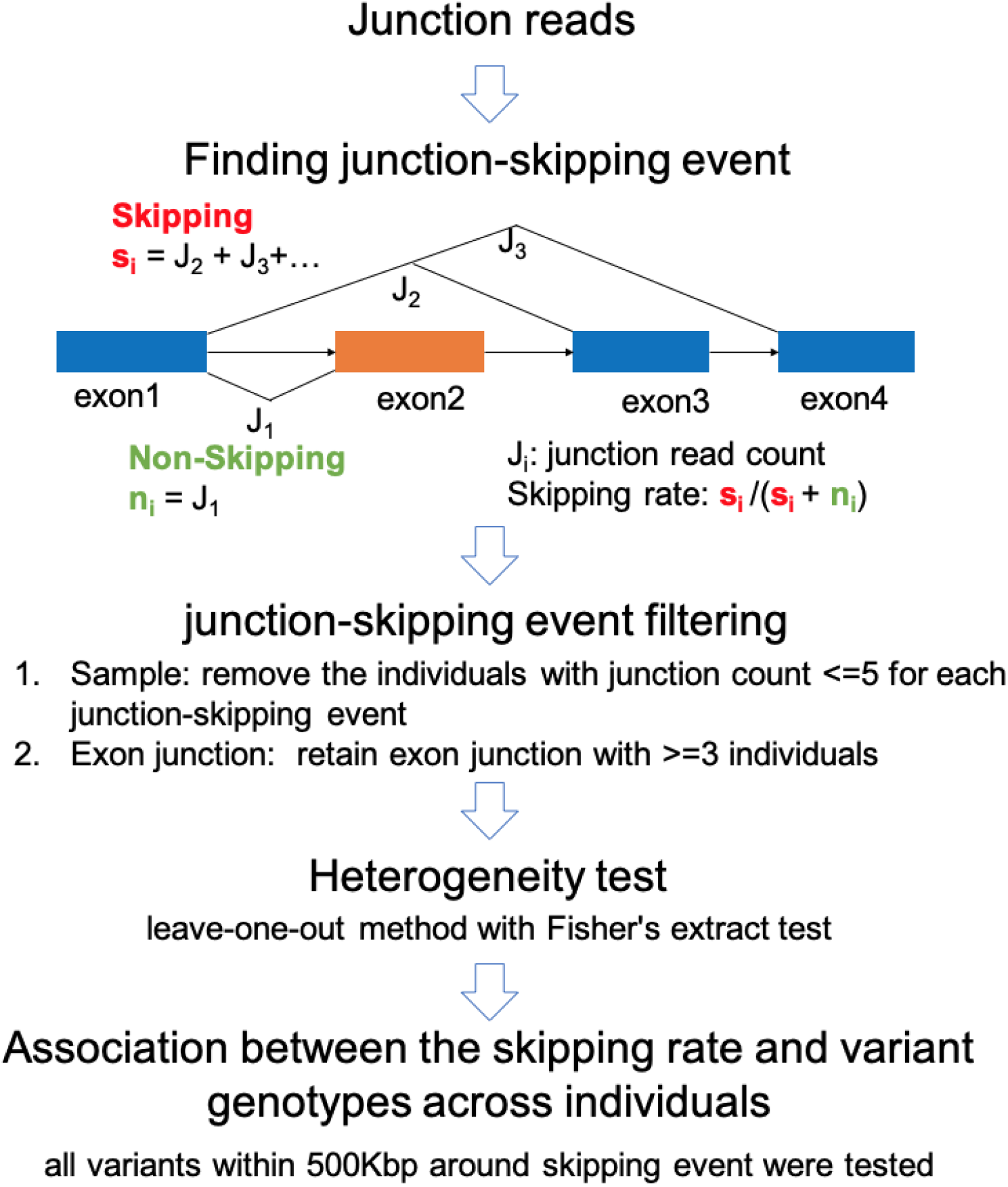
Overview of the jsQTL method. 5’ junction of exon2 is the tested junction. J is junction read count.

To improve the statistical power, we only kept an individual *i* if *n*_*i*_ + *s*_*i*_ > 5. After this, we only kept junctions that have at least 3 individuals remaining. We note that in order to test for variants regulating junction-skipping, we need to observe variations in the junction-skipping pattern in our dataset: if a junction has a homogeneous rate of being skipped across all individuals, it is not possible to test whether the skipping of this junction is regulated by a genomic variant. Therefore, we only included junctions that have heterogeneous skipping rate across samples to find their regulatory variants (Methods), which further reduced the number of tested junctions to 831 - 8,493 per tissue or cell type (Supplementary Table S1).

We tested the association between variants and junction-skipping rate to identify variants regulating junction-skipping (Methods). We did this for common variants with minor allele frequency (MAF) > 0.05 within 500 kilo base-pairs (kb) up and downstream of the junction of interest. We performed 100 permutations to establish the *P*-value threshold corresponding to 0.05 false-positives per genome-wide analysis (Methods). Using this threshold, we found 47 - 574 junction-skipping events significantly regulated by variants, which were fine-mapped to 789 - 12,593 variants in the 95% credible sets (Methods and Supplementary Table S2). 5-130 junction-skipping events were mapped to a single regulatory variant (Supplementary Table S3), and 6 - 154 junction-skipping events were mapped to 2 - 5 candidate variants (Figure 2A). These regulatory variants are shared extensively across tissues: over 50% jsQTLs present in 2+ tissues (Figure 2B). A customizable browser (http://broad.io/jsqtl) is available to review the detailed jsQTL result.

**Figure 2.**
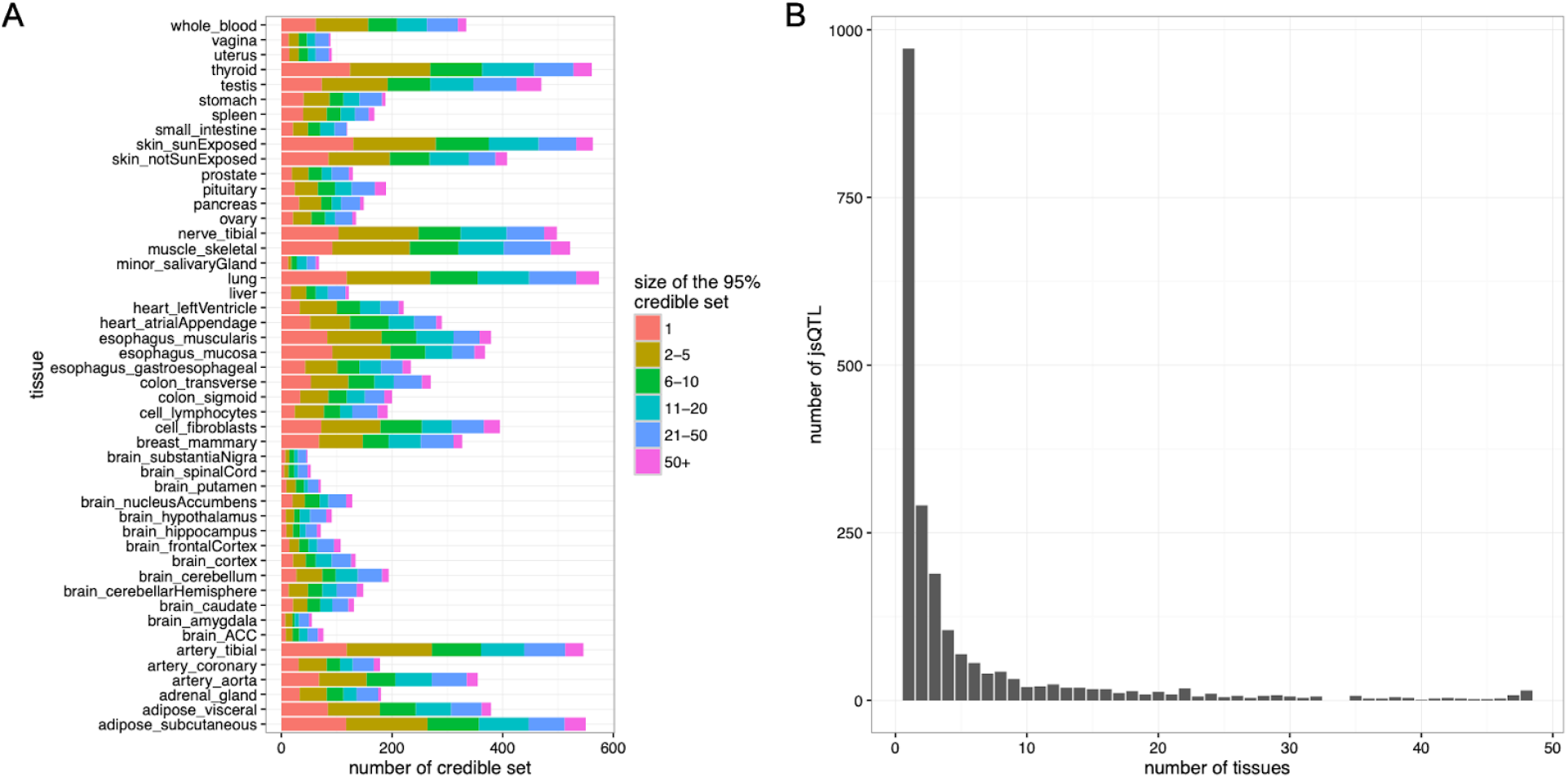
Summary of jsQTL 95% credible set across tissues. a) Number of variants in 95% credible set of each jsQTL, by tissue. b) Number of jsQTLs shared across tissues.

We found that jsQTLs are located close to the skipped junction (Figure 3). To evaluate this quantitatively, for each variant, we took its best probability for being a jsQTL across all tissues and summed up the probabilities across all junctions tested. We found variants within 5,000bp up- and down-stream of the skipped junction account for 46.34% jsQTL probability while only cover 1% (10,000bp / 1,000,000bp) of the tested genomic region. Additionally, we found that 70%-100% of jsQTLs that were mapped to a single variant were within 500bp up- or down-stream of the skipped junction (Supplementary Figure S1).

**Figure 3.**
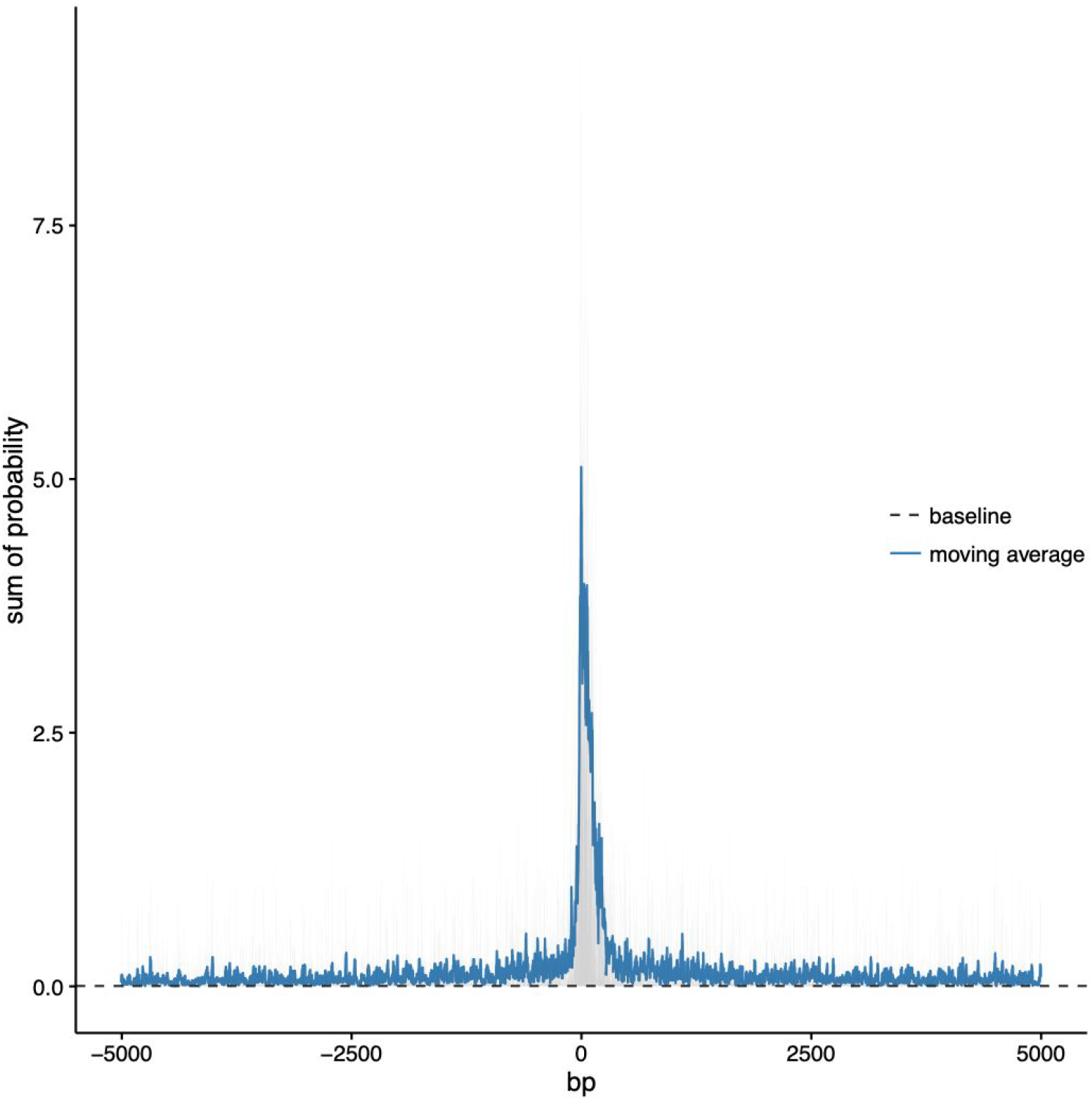
Sum of jsQTL probability for variants close to the skipped junction. Variants within 5,000bp up and down stream of the skipped intron-exon junction were plotted. The position of skipped junction is 0 on the x-axis. The solid line is the moving average of the summed probability calculated with a window size of 10bp. The dashed line is the baseline probability of the full test region (± 500kb), calculated as the average probability across all the variants (Baseline probability is defined as the average posterior probability per base-pair across the tested region).

A few jsQTLs mapped to single variants have been validated in previous studies. For example, we found rs1800693 regulating the skipping of exon 6 in *TNFRSF1A* across 35 tissues (Supplementary Figure S2A and B). Previous studies using *in vitro* minigene splicing assays showed that the G allele of rs1800693 led to the skipping of exon 6^20^. We also found rs74390, 6 nucleotides downstream of the splice acceptor site, regulating the skipping of the 4th exon of *EMID1* across 8 tissues (Supplementary Figure S2C and D). This observation was validated by qRT-PCR in lymphoblastoid cell lines^21^. Lastly, we found rs3795859, located in intron 35 of *LRPPRC*, regulating the skipping of exon 35 across 14 tissues (Supplementary Figure S2E and F). In a minigene assay, the T allele of rs3795859 was shown to increase the exclusion of exon 35 in Hela and HEK293T cell lines^22^.

To investigate the contribution of jsQTLs to human complex traits and disorders, we fine-mapped genome-wide significant loci associated with 23 human complex traits and disorders (Methods and Supplementary Table S4 and S5). We found a handful of jsQTL variants with high posterior probability that also have high posterior probability for being a disease causal variant, suggesting a link between jsQTL and human complex traits and disorders (Supplementary Table S6). More specifically, we found 15 variants across 48 tissues that have posterior probability greater than 10% for both jsQTL and human complex traits/ disorders (Supplementary Table S7). One variant, rs11589479, was found to regulate the skipping of the 19th exon in *ADAM15* in all tested tissues (Supplementary Figure S3A and B). *ADAM15* works as a mediator of mechanisms underlying inflammation and is associated with Crohn’s Disease^23^. In addition, we found rheumatoid arthritis (RA) associated variants (rs2304256 and rs34725611) regulating junction-skipping in *TYK2*, glycated haemoglobin A1c (HbA1c) associated variant (rs267738) regulating junction-skipping in *CERS2* and many height-associated variants regulating junction-skipping in *CDC16, LTBP2*, *ADAMTSL3*, *SEC16A*, *C12orf23*, *NUCB2, GFPT2*, and *SPAG8* across various tissues (Supplementary Figure S3C and D).

Variants may have small posterior probability in fine-mapping due to the limited statistical power or complex linkage disequilibrium (LD) structure. jsQTL analysis may provide additional information to resolve the disease associations. For example, a genetic association with IBD near *SP140*, which is predominantly expressed in immune cells, was mapped to 31 variants^4^. Our jsQTL analysis found rs28445040, one of the 31 credible variants, regulating the skipping of the 7th exon in *SP140.* rs28445040 is the most probable jsQTL variant with 39% posterior probability in lymphocytes (similar results in whole blood and spleen), suggesting that rs28445040 could be an IBD causal variant (Supplementary Figure S3E and F) through regulating the splicing of *SP140*.

On the genome-wide scale (Methods), we noted that jsQTL variants implicate human complex traits and disorders in a tissue specific manner. We found that jsQTLs in whole blood, spleen, lymphocytes, and colon implicate genetic loci associated with inflammatory bowel diseases (IBD), jsQTLs in pancreas implicate genetic loci associated with type 2 diabetes (T2D), and jsQTLs in whole blood and brain tissues (frontal cortex, cerebellum, etc.) implicate schizophrenia genetic loci. Interestingly, genetic associations with height were implicated by jsQTLs in almost all tested tissues and cell lines. We also note that if no connection was observed between jsQTL and disease genetic associations, this could either indicate that the disease/trait is not affected by jsQTL or that the disease association study does not have sufficient power. To reflect this observation, we measured the power of the trait/disease association studies using the variance explained by the fine-mapped variants (Methods), which varies trait by trait from 0.14% to 20.25% (Figure 4A). As the power of the genetic studies increases, the disease-tissue specificity for jsQTL variants becomes more apparent (Figure 4B).

**Figure 4.**
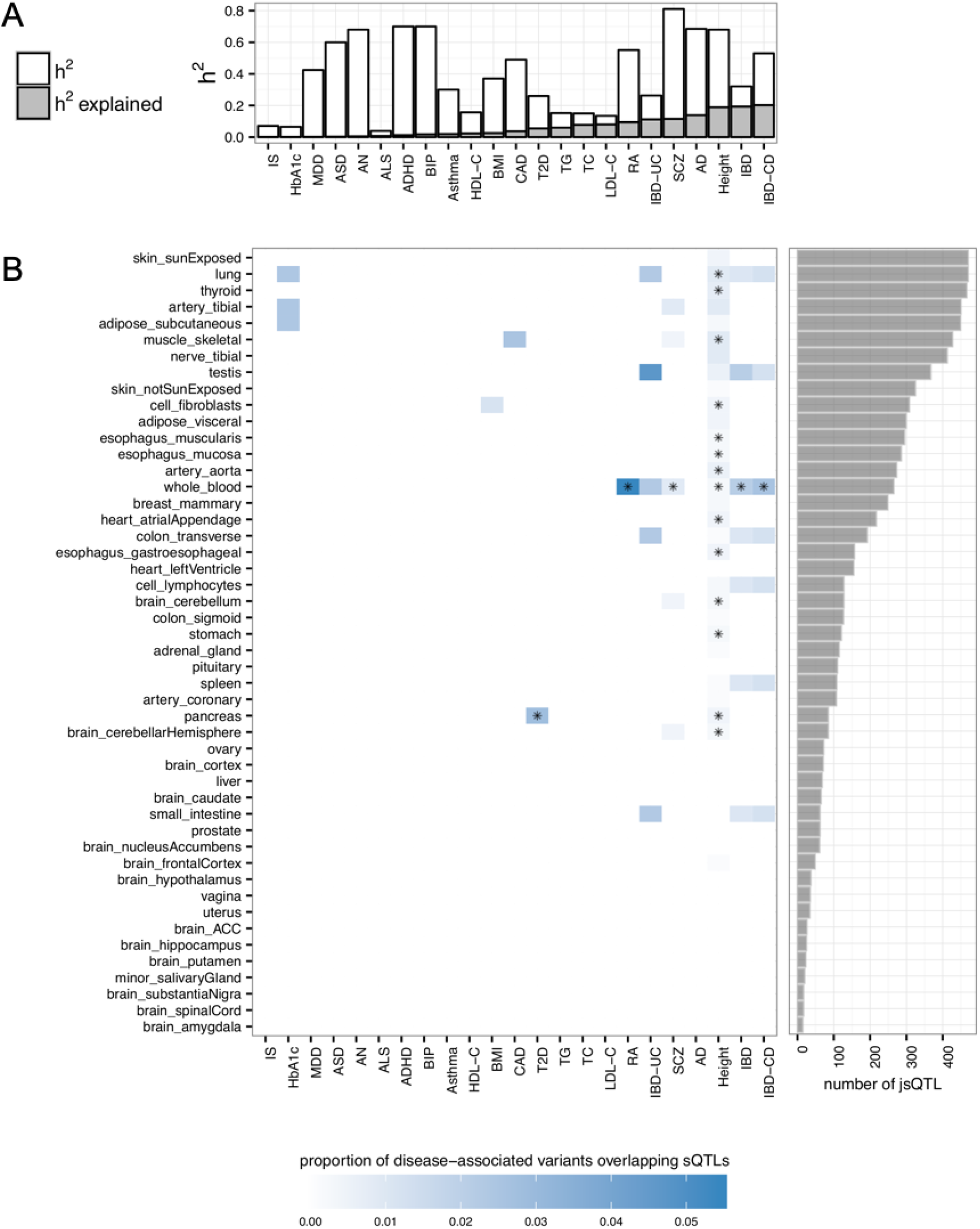
Tissue-specific overlap between jsQTL and disease-associated variants. (A) Common variant-based heritability (white) and the variance explained by fine-mapped variants for each disease/traits(grey). (B) Tissue-specific overlap between jsQTL and disease-associated variants. Color in the heatmap was based on the proportion of disease-associated variants overlapping jsQTLs (Method). * indicates significance beyond the Bonferroni corrected *P*-value threshold (*P*-value < 0.05). TG: Triglycerides, TC: Total cholesterol, T2D: Type 2 diabetes, SCZ: Schizophrenia, RA: Rheumatoid arthritis, MDD: Major Depressive Disorder, LDL-C: low-density lipoprotein cholesterol, IS: Ischemic stroke, IBD-UC: ulcerative colitis, IBD-CD: Crohn’s Disease, IBD: Inflammatory Bowel Disease, Height: height, HDL-C: high-density lipoprotein cholesterol, HbA1c: Glycated hemoglobin, CAD: Coronary artery disease, BMI: Body Mass Index, BIP: Bipolar disorder, Asthma: Asthma, ASD: Autism, AN: Anorexia nervosa, AD: Alzheimer’s disease, ALS: amyotrophic lateral sclerosis, ADHD: Attention-deficit/hyperactivity disorder

## Discussion

In this study, we developed an approach to identify junction-skipping regulatory variants using data from GTEx and investigated their connections with human complex disorders. Alternative splicing is a complex event which had been characterized using transcript- or exon/intron-focused methods. We proposed a new junction-focused method with reduced complexity, providing a fresh look into alternative splicing in human transcriptome. This method is able to detect alternative splicing regardless of the number of exons skipped, although it is unable to provide details regarding individual exons.

We performed fine-mapping to estimate the causal probability for variants regulating the junction-skipping. We found that splicing regulatory variants clearly contribute to human complex disorders and traits in a tissue specific manner, providing a valuable resource for functionally characterizing the role of the noncoding genome in complex diseases and traits. Similarly, we fine-mapped disease genetic associations to a smaller and precise subset of variants. By only focusing on associations that can be mapped to a small set of variants, we achieved better specificity and reduced spurious colocalizations. There is clear evidence that the majority of variants identified in GWAS contribute to the disease risk through their impact on gene regulatory functions in specific tissues^24^. Findings in this study agree with previous findings that splicing QTL are major contributors to complex diseases/traits^8,14^. For example, researchers found splicing QTLs in LCLs enriched in variants associated with autoimmune-disease and splicing QTLs in human brains enriched with variants associated with schizophrenia^8,14,25^. In this study, we also found T2D-associated variants enriched in jsQTLs from the pancreas, the organ producing insulin; and variants associated with IBD and Rheumatoid arthritis (RA) enriched in jsQTLs from the immune-related cell lines and tissues including whole blood, spleen, and fibroblasts. These findings are supported by previous reports that genetic associations with CD co-localize with immune-cell chromatin peaks^4^, and that the CD is an autoimmune disorder resulting from an impaired innate immunity or an overactive Th1 and Th17 cytokine response^26,27^.

One limitation of this study is the sample size. Sample size plays an important role in association and fine-mapping studies^4,28^. As the sample size increases, more jsQTL can be discovered (Supplementary Figure S4), showing that we have not yet saturated the power and more samples are needed to complete the allelic spectrum of splicing regulatory variants. Future studies such as GTEx release v8 will deliver additional samples to further our understanding of the connection between jsQTL and complex diseases.

This study only investigated common variants with MAF > 5% in the genome as the power for testing less frequent variants is low. The frequencies for variants at the 5’ or 3’ essential splicing sites tend to be low because they are often under purifying selection. Therefore, most of the regulatory variants we identified in this study were not at the 5’ or 3’ splice sites and do not have full penetrance. Rather, they have a moderate effect on the exon splicing and affect the splicing efficiency. This is consistent with the view that mutations in other positions of the consensus sequence that decrease splice site strength can partially or completely inhibit usage of the splice site^29^.

## Methods

### Dataset

The RNA-seq and genotype data are from GTEx release v7 (Supplementary Table S1). The GWAS summary statistics used for fine-mapping were downloaded from published studies (Supplementary Table S4).

### Identification of Junction-Skipping events

To identify the junction-skipping events, uniquely mapped junction reads were extracted from the GTEx alignment bam file using samtools^30^ and regtools (https://regtools.readthedocs.io/en/latest/). The uniquely mapped junction reads in samples from the same tissue were pooled to identify junction-skipping event from the canonical transcripts annotation (APPRIS) ^31^. For each tested exon-exon junction, we counted the number of non-skipping junction reads as *n*_*i*_ and the number of skipping junction reads as *s*_*i*_, with *i* denoting the individual *i*. As shown in Figure 1, non-skipping junction reads are the reads covering exons 1 and 2 in the canonical transcript; and skipping junction reads are the reads covering exon 1 and another exon downstream of exon 2 (including 3, 4, …) in the canonical transcript. As discussed in the Result, for each junction-skipping event, samples with no more than 5 junction reads (*n*_*i*_ + *s*_*i*_≤5) were removed. In addition, each junction-skipping event with less than 3 samples was not tested due to low statistical power.

### Heterogeneity test for outliers

To reduce the multiple testing burden, we selected a subset of junction-skipping events that are heterogeneous across individuals. This is because the events that are homogeneous across all individuals have no power for association tests. We used the leave-one-out method to test the heterogeneity. For a tissue and a junction-skipping event, assume there are *m* samples and *j* is the sample being tested for heterogeneity. We first calculated the number of skipped junction read counts across all samples except for sample *j* as 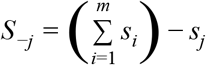. Similarly, the sum of the non-skipped junction read counts without sample *j* is 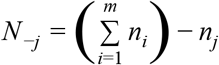. We then applied Fisher’s exact test on sample *j* using the 2-by-2 matrix: 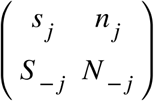.

We denote the *P*-value from the Fisher’s exact test as *P*_*j*_, and the Bonferroni corrected *P*-value can be calculated as *P*′_*j*_ = 1 − (1 − *P*_*j*_)^*m*^. We defined “the number of outliers” as 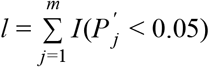, in which *I*() is Indicator function with a value of 1 if *P*′_*j*_ < 0.05 and value of 0 otherwise. We only test the junction-skipping events that have *l* > 5.

### Association analysis between junction-skipping and variants

For each variant, junction-skipping event and tissue, we divide samples into two groups (*G1 and G2*) based on their genotypes:

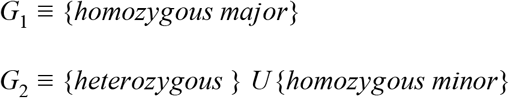

We then calculate the skipping rate for each individual and group them by:

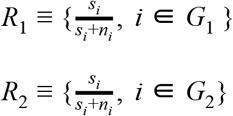

We test the difference of the skipping rates between the two groups (*R*1 *v.s*. *R*2) using the Mann-Whitney-Wilcoxon test, and denote the result as *P*_*w*_. Variants with minor allele frequency ≥ 5% and within ±500 kb of the skipped intro-exon junction were tested for their associations with the junction-skipping event.

### Establish the genome-wide significance threshold

Variants are correlated because of LD so the number of independent tests can be smaller than the number of exon-variant pairs tested. We used a permutation test to estimate the test burden and the *P*-value threshold after multiple testing corrections. We performed the test on skeletal muscle because it has the largest sample size and therefore, more exon junction tested due to the power (a conservative choice). For skeletal muscle samples, we randomly shuffled the sample ID 100 times, for each shuffle, we used the shuffled ID to connect the skipping rate and the genotypes, and performed association tests for all exon-variant pairs. We noted a well-calibrated *P*-value distribution from the permutation test (Supplementary Figure S5). The minimum *P*-value across all tested exon-variant pairs in each shuffle was recorded for a list of 100 “minimum” *P*-values. The 5th smallest *P*-value in the list, which was 5×10^−10^, was used as the genome-wide significant *P*-value threshold (corresponding to 5% genome-wide false-positive rate).

### Fine-mapping of jsQTL signals

We fine-mapped all junction-skipping events that have at least one variant with MAF ≥ 5% significantly associated (*P*-value < 5×10^−10^). We used a published fine-mapping method^4^ with default parameters (R2< 0.4), and LD calculated from individual-level genotypes from GTEx. We calculated the causal posterior probability for each variant in the region, and constructed the smallest set of variants that contain the causal variant with 95% confidence (95% credible set). The extended MHC region (chr6 22.5M - 33.5M) was not included because of its long-range LD.

### Fine-mapping of complex disease loci

We took the fine-mapping results from a published fine-mapping study^4^ for the inflammatory bowel diseases. For other diseases and traits, we used the same fine-mapping approach we used for jsQTL^4^. We downloaded the summary statistics from 23 GWAS (Supplementary Table S4). For each disease or traits, we fine-mapped genetic associations that reached genome-wide significance (*P*-value < 5×10^−8^) and have MAF ≥ 5%. Variants with R^2^ < 0.4 and are within 250 kb up- and down-stream of the most significant variant from GWAS were used in fine-mapping. We used PLINK^32^ to compute the pairwise R^2^ between the most significant GWAS variant and other variants in each locus using the 1000 Genomes Reference Panel Phase 3 European (EUR) population^33^. We only fine-mapped the primary association in each locus. The outcome from the fine-mapping analysis was a set of variants that contain the causal variant with 95% confidence (95% credible set) with their causal posterior probability. The extended MHC region (chr6 22.5M - 33.5M) was not included because of its long-range LD.

### Overlap between jsQTL variants and variants associated with complex diseases and traits

To characterize the contribution of jsQTLs to human complex diseases and traits, we first counted how many disease associations overlap the jsQTL associations in each tissue. An overlap is defined as the 95% credible sets for both associations share at least one variant. This number was then normalized by the total number of genetic associations fine-mapped for the disease, representing the proportion of the disease associations regulating junction-skipping. We then used permutations to establish the distribution of this proportion under the null, and use the distribution to evaluate the statistical significance of the observed proportion. The permutation was performed 10,000 times for each tissue-disease pair by shifting variants within 95% credible sets for each disease at a random direction for a random distance between 0 and 500 kb. This permutation was performed circularly meaning that variants at the end of the locus will be moved to the start of the locus. *P*-value was calculated as the number of times the proportion from the permuted data is greater than the original proportion in the unpermuted data, divided by the number of permutations performed (10,000). The *P*-value significance threshold was corrected for the number of disease-tissue pairs with at least one overlap in the unpermuted data: 0.05 / 60=0.0008. To improve the signal-to-noise ratio, we only included jsQTLs that showed reasonable tissue specificity (detected in < 35 out of 48 tissues, Figure 2) and only disease and tissues credible sets that were mapped to ≤50 variants.

### Variance explained by fine-mapped disease associations and the disease heritability

We calculated the variance explained by the fine-mapped disease associations using the variant with the largest posterior probability in each credible set as the proxy. The variance explained for each variant was then calculated as 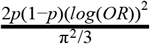 for categorical traits and 2*p*(1 − *p*)(*beta*)^2^ for quantitative traits ^34^, where p is minor allele frequency in the 1000 Genomes Reference Panel Phase 3 European (EUR) population, OR is odds ratio, and beta is the per allele effect on the quantitative trait. The variance explained by all credible sets for a disorder is then the sum of variance explained by each proxy variant. The heritability for a trait/disease was taken from SNPedia^35^ and LD Hub (if not available in SNPedia)^36^.

## Acknowledges

We acknowledges NIDDK K01DK114379 and funding support from Stanley Center for Psychiatric Research.

## Supplement Tables

1. Supplementary Table S1. Sample information for jsQTL analysis and summary results
2. Supplementary Table S2. jsQTL variants mapped to 95% credible sets
3. Supplementary Table S3. jsQTLs mapped to single variant
4. Supplementary Table S4. Sample information for traits/diseases fine-mapping analysis and summary results
5. Supplementary Table S5. Human traits/disease associated variants fine-mapped to 95% credible sets
6. Supplementary Table S6. Variants fine-mapped to 95% credible sets for both jsQTL and human complex traits/disorders
7. Supplementary Table S7. Variants with posterior probability > 10% for both jsQTL and human complex traits/ disorders

## Supplement Figures

1. Supplementary Figure S1. Distribution of distance between the skipped exon and its jsQTLs.
2. Supplementary Figure S2. Examples of jsQTLs mapped to a single variant (labeled)
3. Supplementary Figure S3. Examples of jsQTLs for disease-associated genes.
4. Supplementary Figure S4. The number of identified jsQTLs increases with the sample size.
5. Supplementary Figure S5. Quantile-quantile (QQ) plot of P-value from the permuted data in skeletal muscle.

## Reference

1. Baralle, F. E. & Giudice, J. Alternative splicing as a regulator of development and tissue identity. Nat. Rev. Mol. Cell Biol. 18, 437–451 (2017).

2. Kelemen, O. et al. Function of alternative splicing. Gene 514, 1–30 (2013).

3. Park, E., Pan, Z., Zhang, Z., Lin, L. & Xing, Y. The Expanding Landscape of Alternative Splicing Variation in Human Populations. Am. J. Hum. Genet. 102, 11–26 (2018).

4. Huang, H. et al. Fine-mapping inflammatory bowel disease loci to single-variant resolution. Nature 547, 173–178 (2017).

5. Gallagher, M. D. & Chen-Plotkin, A. S. The Post-GWAS Era: From Association to Function. Am. J. Hum. Genet. 102, 717–730 (2018).

6. Montes, M., Sanford, B. L., Comiskey, D. F. & Chandler, D. S. RNA Splicing and Disease: Animal Models to Therapies. Trends Genet. 35, 68–87 (2019).

7. Wang, G.-S. & Cooper, T. A. Splicing in disease: disruption of the splicing code and the decoding machinery. Nat. Rev. Genet. 8, 749–761 (2007).

8. Li, Y. I. et al. RNA splicing is a primary link between genetic variation and disease. Science 352, 600–604 (2016).

9. Pagani, F. & Baralle, F. E. Genomic variants in exons and introns: identifying the splicing spoilers. Nat. Rev. Genet. 5, 389–396 (2004).

10. Zheng, S. & Chen, L. A hierarchical Bayesian model for comparing transcriptomes at the individual transcript isoform level. Nucleic Acids Res. 37, e75 (2009).

11. Conesa, A. et al. A survey of best practices for RNA-seq data analysis. Genome Biol. 17, 13 (2016).

12. Anders, S., Reyes, A. & Huber, W. Detecting differential usage of exons from RNA-seq data. Genome Res. 22, 2008–2017 (2012).

13. Shen, S. et al. rMATS: robust and flexible detection of differential alternative splicing from replicate RNA-Seq data. Proc. Natl. Acad. Sci. U. S. A. 111, E5593–601 (2014).

14. Li, Y. I. et al. Annotation-free quantification of RNA splicing using LeafCutter. Nat. Genet. 50, 151–158 (2018).

15. Vaquero-Garcia, J. et al. A new view of transcriptome complexity and regulation through the lens of local splicing variations. Elife 5, e11752 (2016).

16. Rivas, M. A. et al. Deep resequencing of GWAS loci identifies independent rare variants associated with inflammatory bowel disease. Nat. Genet. 43, 1066–1073 (2011).

17. Malik, M. et al. CD33 Alzheimer’s risk-altering polymorphism, CD33 expression, and exon 2 splicing. J. Neurosci. 33, 13320–13325 (2013).

18. Matesanz, F. et al. A functional variant that affects exon-skipping and protein expression of SP140 as genetic mechanism predisposing to multiple sclerosis. Hum. Mol. Genet. 24, 5619–5627 (2015).

19. GTEx Consortium. Human genomics. The Genotype-Tissue Expression (GTEx) pilot analysis: multitissue gene regulation in humans. Science 348, 648–660 (2015).

20. Gregory, A. P. et al. TNF receptor 1 genetic risk mirrors outcome of anti-TNF therapy in multiple sclerosis. Nature 488, 508–511 (2012).

21. Mucaki, E. J. & Rogan, P. K. Expression changes confirm predicted single nucleotide variants affecting mRNA splicing. bioRxiv 549089 (2019). doi:10.1101/549089

22. Gaweda-Walerych, K., Mohagheghi, F., Zekanowski, C. & Buratti, E. Parkinson’s disease-related gene variants influence pre-mRNA splicing processes. Neurobiol. Aging 47, 127–138 (2016).

23. Momozawa, Y. et al. IBD risk loci are enriched in multigenic regulatory modules encompassing putative causative genes. Nat. Commun. 9, 2427 (2018).

24. Hormozdiari, F. et al. Leveraging molecular quantitative trait loci to understand the genetic architecture of diseases and complex traits. Nat. Genet. 50, 1041–1047 (2018).

25. Takata, A., Matsumoto, N. & Kato, T. Genome-wide identification of splicing QTLs in the human brain and their enrichment among schizophrenia-associated loci. Nat. Commun. 8, 14519 (2017).

26. Marks, D. J. B. & Segal, A. W. Innate immunity in inflammatory bowel disease: a disease hypothesis. J. Pathol. 214, 260–266 (2008).

27. Elson, C. O. et al. Monoclonal anti-interleukin 23 reverses active colitis in a T cell-mediated model in mice. Gastroenterology 132, 2359–2370 (2007).

28. Visscher, P. M. et al. 10 Years of GWAS Discovery: Biology, Function, and Translation. Am. J. Hum. Genet. 101, 5–22 (2017).

29. Cieply, B. & Carstens, R. P. Functional roles of alternative splicing factors in human disease. Wiley Interdiscip. Rev. RNA 6, 311–326 (2015).

30. Li, H. et al. The Sequence Alignment/Map format and SAMtools. Bioinformatics 25, 2078–2079 (2009).

31. Rodriguez, J. M. et al. APPRIS: annotation of principal and alternative splice isoforms. Nucleic Acids Res. 41, D110–7 (2013).

32. Purcell, S. et al. PLINK: a tool set for whole-genome association and population-based linkage analyses. Am. J. Hum. Genet. 81, 559–575 (2007).

33. 1000 Genomes Project Consortium et al. A global reference for human genetic variation. Nature 526, 68–74 (2015).

34. Pawitan, Y., Seng, K. C. & Magnusson, P. K. E. How many genetic variants remain to be discovered. PLoS One 4, e7969 (2009).

35. Cariaso, M. & Lennon, G. SNPedia: a wiki supporting personal genome annotation, interpretation and analysis. Nucleic Acids Res. 40, D1308–12 (2012).

36. Zheng, J. et al. LD Hub: a centralized database and web interface to perform LD score regression that maximizes the potential of summary level GWAS data for SNP heritability and genetic correlation analysis. Bioinformatics 33, 272–279 (2017).

