## Supplementary Figures for "Junction-skipping regulation in complex disease"

**Supplementary Figure S1**

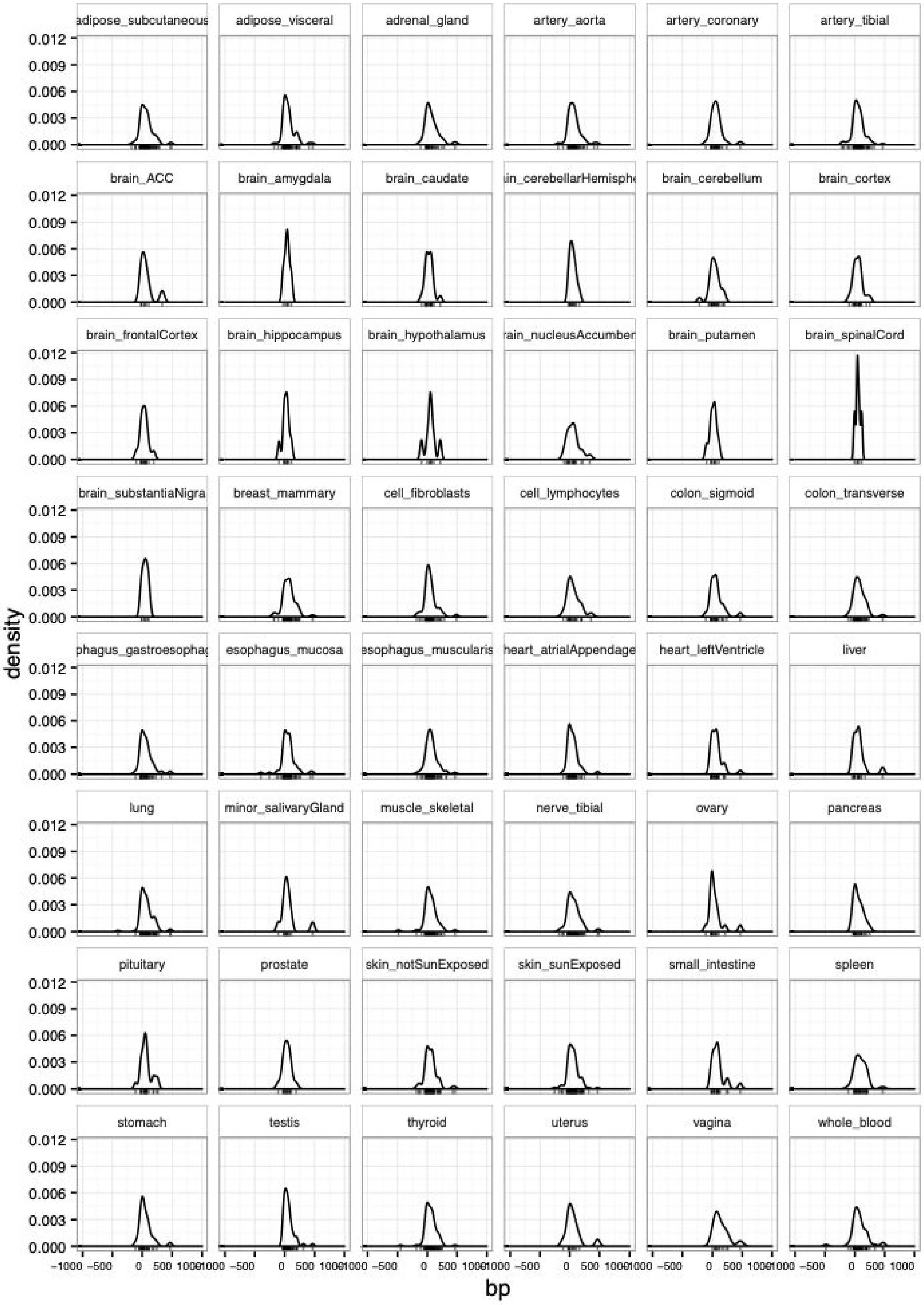

**Supplementary Figure S1.** Distribution of distance between the skipped exon and its sjQTLs. Only sjQTLs mapped to single variant is used in this figure. 0 on the x-axis is the position of skipped junction.

**Supplementary Figure S2**

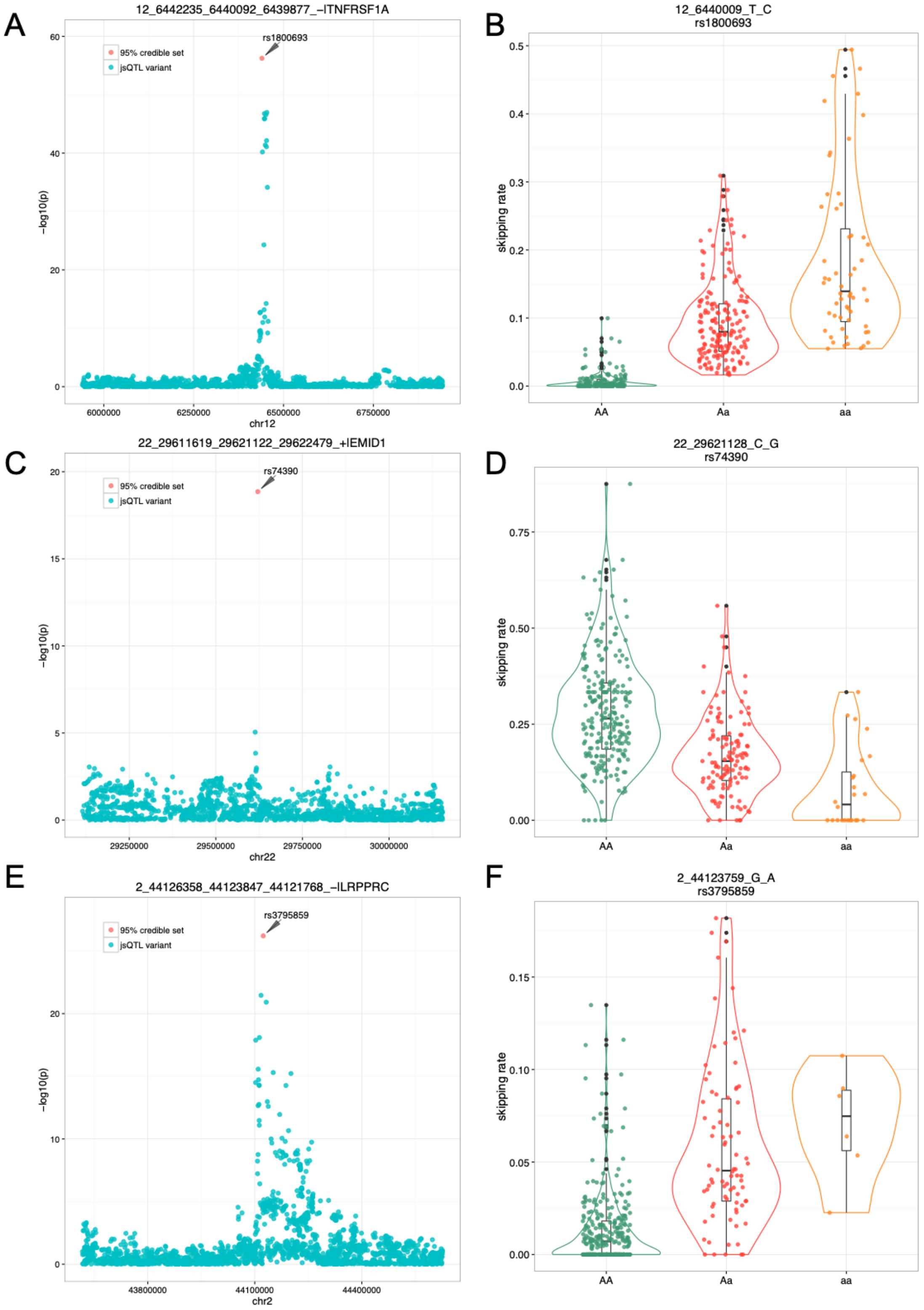


**Supplementary Figure S2.** Examples of jsQTLs mapped to a single variant (labeled). A and B) rs1800693 regulating the skipping of exon 6 in *TNFRSF1A* in lung. C and D) rs74390 regulating the skipping of exon 4 in *EMID1* in lung. E and F) rs3795859 regulating exon 35 in *LRPPRC* in lung.

**Supplementary Figure S3**

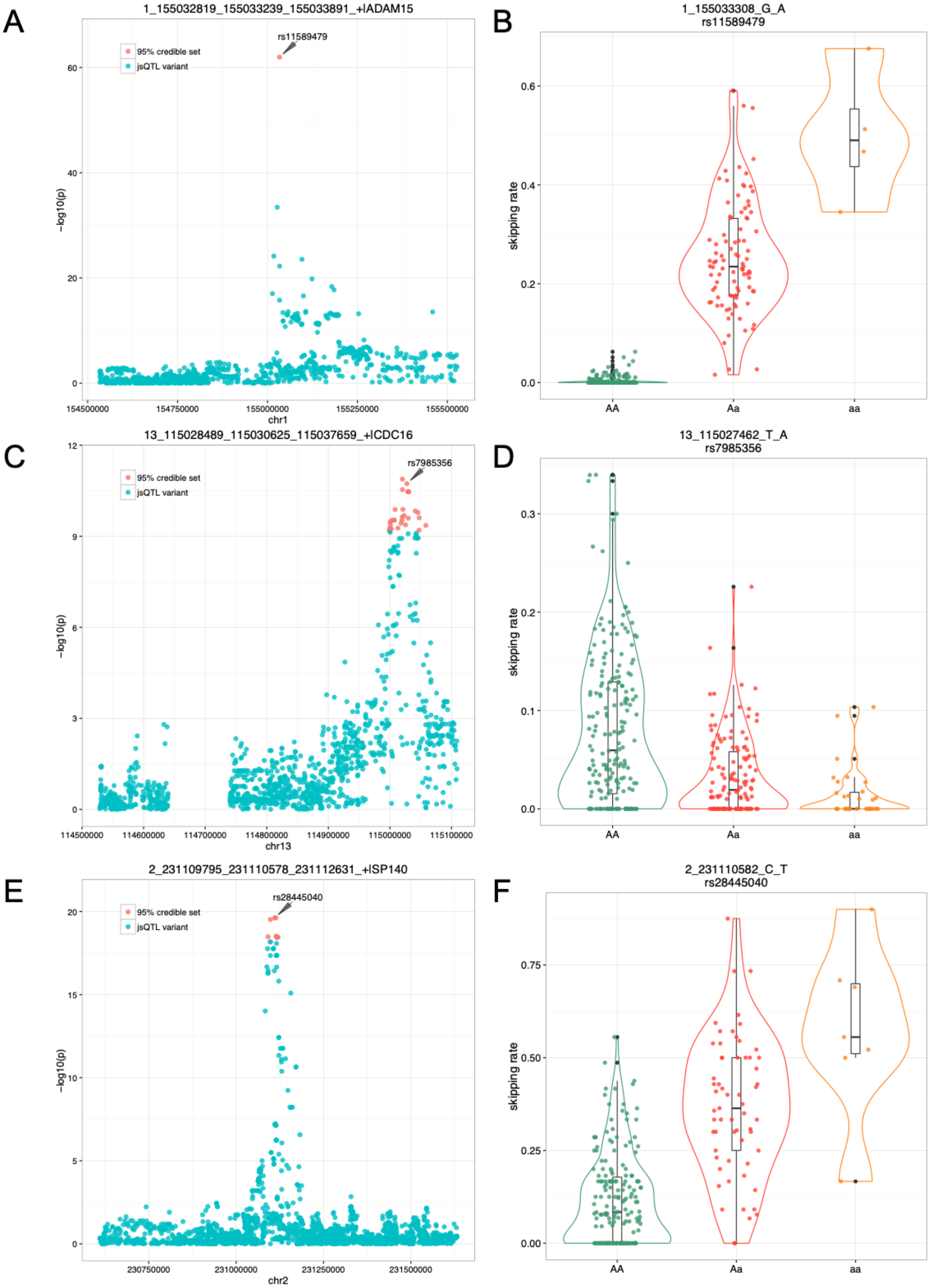


**Supplementary Figure S3.** Examples of jsQTLs for disease-associated genes. A and B) rs11589479 regulating the skipping of *ADAM15* exon 19 and is associated with IBD and CD. C and D) rs7985356 regulating the skipping of *CDC16* exon 17 and is associated with height. E and F) rs28445040 regulating the skipping of *SP140* exon 7 and is associated with IBD and CD.

**Supplementary Figure S4**

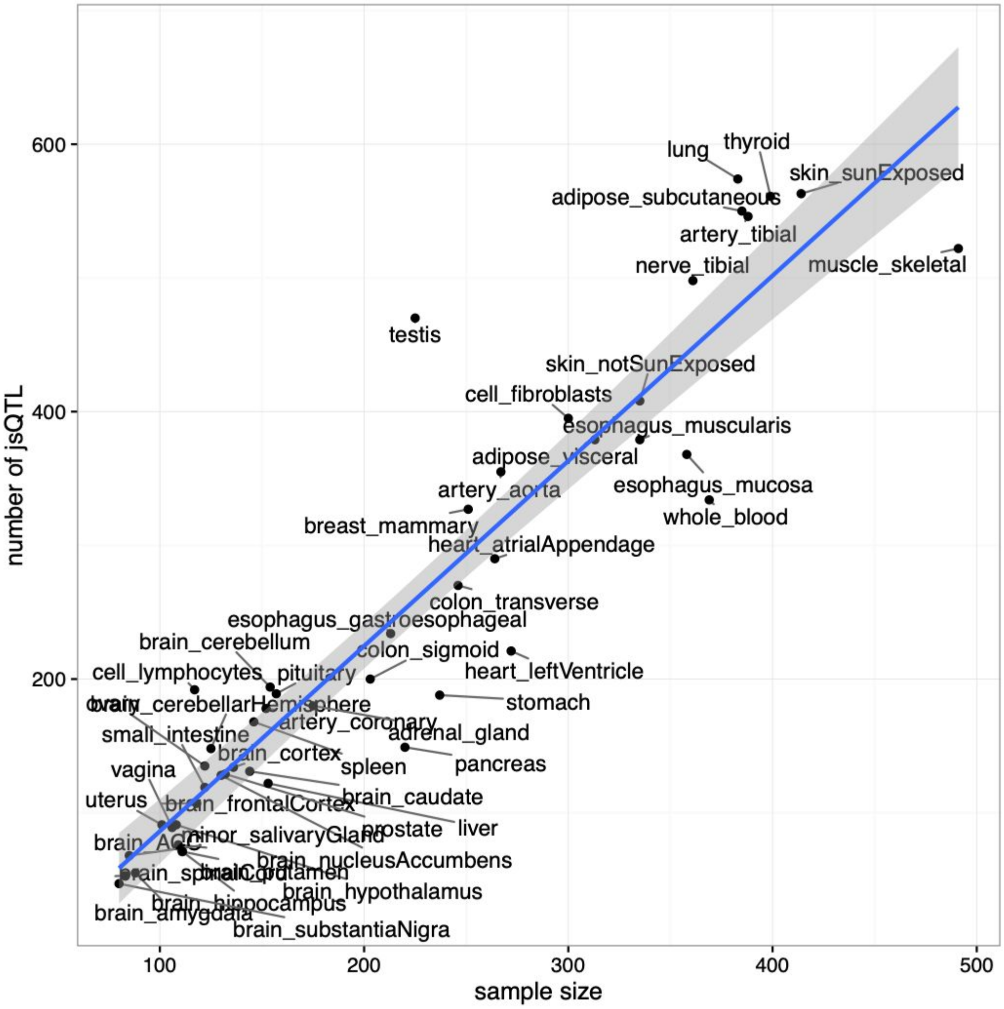


**Supplementary Figure S4.** The number of identified jsQTLs increases with the sample size.

**Supplementary Figure S5**

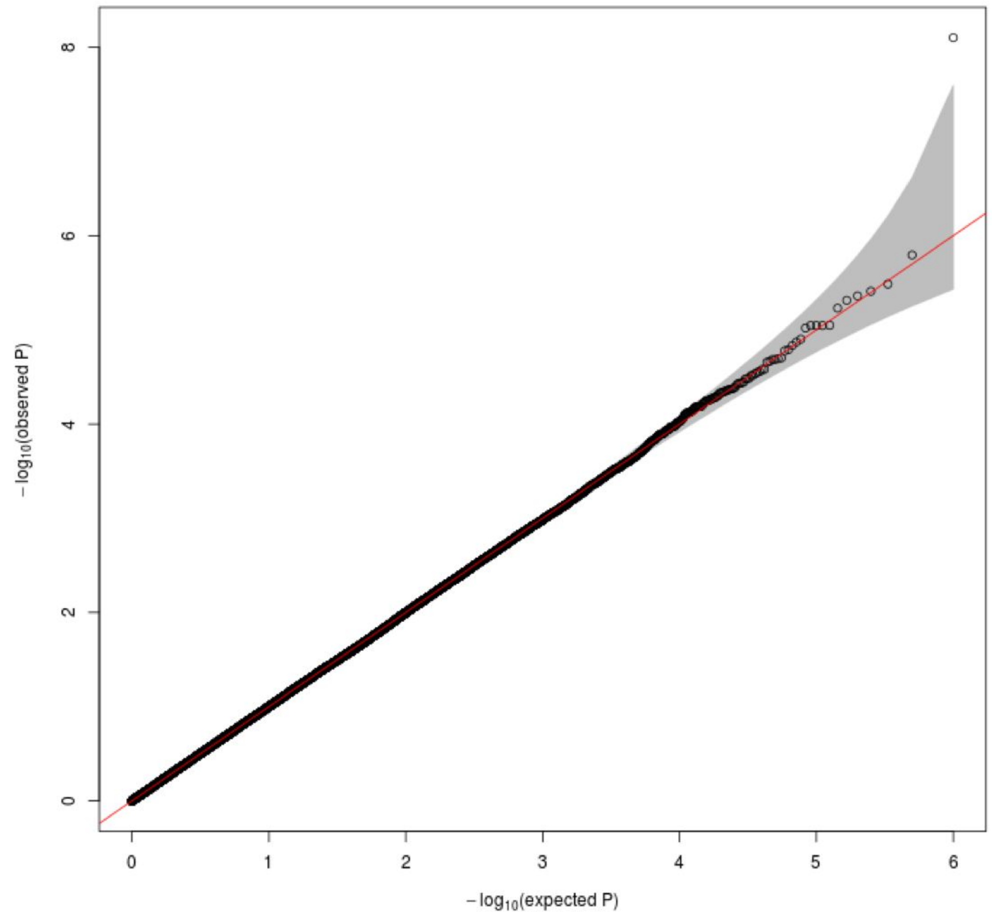


**Supplementary Figure S5**. Quantile-quantile (QQ) plot of *P*-value from the permuted data in skeletal muscle. One million *P*-values were random samples for plotting the QQ plot.
